# Mapping eQTL by leveraging multiple tissues and DNA methylation

**DOI:** 10.1101/069534

**Authors:** Chaitanya R. Acharya, Kouros Owzar, Andrew S. Allen

## Abstract

**Background:** DNA methylation is an important tissue-specific epigenetic event that influences transcriptional regulation of gene expression. Differentially methylated CpG sites may act as mediators between genetic variation and gene expression, and this relationship can be exploited while mapping multi-tissue expression quantitative trait loci (eQTL). Current multi-tissue eQTL mapping techniques are limited to only exploiting gene expression patterns across multiple tissues either in a joint tissue or tissue-by-tissue frameworks. We present a new statistical approach that enables us to model the effect of germ-line variation on tissue-specific gene expression in the presence of effects due to DNA methylation.

**Results:** Our method efficiently models genetic and epigenetic variation to identify genomic regions of interest containing combinations of mRNA transcripts, CpG sites, and SNPs by jointly testing for genotypic effect and higher order interaction effects between genotype, methylation and tissues. We demonstrate using Monte Carlo simulations that our approach, in the presence of both genetic and DNA methylation effects, gives an improved performance (in terms of statistical power) to detect eQTLs over the current eQTL mapping approaches. When applied to an array-based dataset from 150 neuropathologically normal adult human brains, our method identifies eQTLs that were undetected using standard tissue-by-tissue or joint tissue eQTL mapping techniques. As an example, our method identifies eQTLs in a BAX inhibiting gene (TMBIM1), which may have a role in the pathogenesis of Alzheimer disease.

**Conclusions:** Our score test-based approach does not need parameter estimation under the alternative hypothesis. As a result, our model parameters are estimated only once for each mRNA - CpG pair. Our model specifically studies the effects of non-coding regions of DNA (in this case, CpG sites) on mapping eQTLs. However, we can easily model micro-RNAs instead of CpG sites to study the effects of post-transcriptional events in mapping eQTL. Our model’s flexible framework also allows us to investigate other genomic events such as alternative gene splicing by extending our model to include gene isoform-specific data.

## Background

It has been long established that regulatory regions in higher eukaryotes activate gene transcription in a tissue-specific manner [1, 2]. These regulatory regions, which affect the binding affinities of transcription factors, are susceptible to both genetic variation and epigenetic modifications that play a coordinated role in regulating tissue-specific gene expression [3, 4, 5, 6, 7]. One form of epigenetic variation is DNA methylation that targets nonmethylated and noncoding GC-rich and CpG-rich regions of the DNA sequence, which constitute approximately 70% of all annotated promoters [8]. DNA methylation is linked to transcriptional silencing, and many CpG island promoters are active in a tissue-specific manner. Previous studies have shown that inter-individual variation in DNA methylation at distinct CpG sites has been consistently linked to genetic variation such as single nucleotide polymorphisms (SNPs), known as methylation eQTLs (mQTLs) [9, 10, 11]. Since an increased DNA methylation at any of the distinct CpG sites located in the promoter regions necessitate chromatin remodeling and subsequent decrease in gene expression, any DNA sequence variation within the CpG-rich regions that disrupts the methylation process may have an opposite effect on gene expression.

Even though, mechanisms which regulate DNA methylation are unclear, it is clear that there is some association between genetic variation and quantitative changes in methylation levels [14]. For example, Catechol-O-methyltransferase (COMT) gene, which is implicated in schizophrenia has a SNP, *Val*^158^ *Met* (rs4680) that is associated with differential COMT expression across regions of the brain during the course of the illness [15]. More specifically, the substitution of a methionine (Met) for a valine (Val) at position 158 results in reduced activity of the COMT enzyme due to reduced protein stability. Methylation of CpG islands associated with the aforementioned variant affect the region-specific expression of COMT [15]. Identifying and studying the mechanisms through which genetic variation, DNA methylation and gene expression interact may provide us yet another clue to understanding regions within the genome that are associated with complex disease phenotypes (Figure 1).

**Figure 1:**
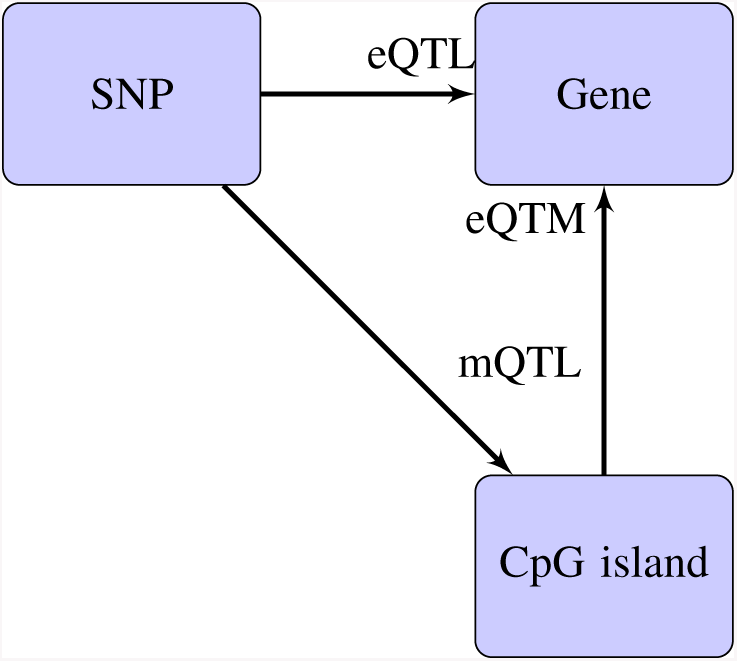
Tissue-specific gene expression is controlled by genetic, epigenetic and transcriptional regulatory mechanisms. Genetic control of gene expression can be defined in terms of SNPs and their associations with gene expression. Expression quantitative trait loci (eQTL) are correlations between SNPs and gene expression. Similarly, epigenetic control of gene expression can be defined in terms of CpG island methylation and their interactions with SNPs. Expression quantitative trait methylations (eQTM) are correlations between gene expression and methylation while methylation quantitative trait loci (mQTL) are correlations between SNPs and methylation. Gene expression can be modeled as a function of SNP and CpG islands.

Current approaches to delineate the role played by both genetic and epigenetic variation in gene expression are limited to identifying statistically significant pairs of mRNA - SNPs and CpG - SNPs by performing independent eQTL and mQTL analyses, respectively, within a tissue-by-tissue (TBT) framework [4, 11, 16]. These pairs are then expanded to combinations of mRNA transcript, CpG site and a SNP wherever the SNP was significantly correlated with either mRNA or CpG site of the mRNA - CpG pair. First, any such TBT analyses have been shown to fall short in fully exploiting patterns across the tissues thus impacting eQTL or mQTL discovery [17, 18, 19]. Second, independent eQTL and mQTL analyses do not reveal any underlying effects of genetic variation on tissue-specific gene expression due to DNA methylation. Consequently, we propose to map eQTLs by leveraging DNA methylation and testing for any higher order interactions among methylation, genotype and tissues. We have previously proposed a score test-based approach to map multi-tissue eQTLs where we model tissue-specificity as a random effect and investigated an overall shift in the gene expression combined with tissue-specific effects due to genetic variants [19]. We extend this framework to include methylation-specific effects and model the combined effect of genetic and epigenetic variation on gene expression.

We show using Monte Carlo simulations that our joint score test is more powerful in teasing out eQTLs by controlling for methylation than any TBT approach that uses methylation as a covariate (TBTm-eQTL). We also show that the new joint score test is better at identifying eQTLs in the presence of DNA methylation than our previously proposed multi-tissue eQTL and TBT methods. Finally, we show that in cases where the interaction effects of DNA methylation are absent, our approach remains competetive. We demonstrate the effectiveness of our method by applying it to a publicly available expression, methylation and SNP array datasets from adult normal brains [4] and show that by jointly analyzing multiple brain regions (tissues), we identify eQTL that may otherwise be not identified by multi-tissue eQTL methods.

## Methods

### Our model

For a given mRNA transcript, tissue-specific gene expression is modeled as a function of genotype and methylation –

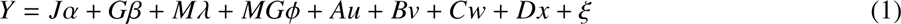

where *Y* is *nt*-dimensional vector of expression levels in *t* tissues and *n* individuals, *α* is a vector of tissue-specific intercepts, *G* is *nt*-dimensional vector of genotypes, β is a fixed effect of genotype across tissue, *M* is *nt*-dimensional vector of methylation levels, λ is an overall methylation-specific fixed effect, *MG* is *nt*-dimensional vector of the product of methylation and genotype, φ is the regression coefficient for genotype and methylation interaction (fixed effect), *u* ~ *N* (0, τ *AA^T^*) is a vector of subject-specific random effect, *v* ~ *N*(0, *γBB^T^*) is a vector of tissue-specific random effects, *w* ~ *N(*0, *δCC^T^*)is a vector of tissue-specific random effects that describe the interaction effect between genotype and methylation is a vector of random effects describing the interaction between genotype, methylation and tissue, 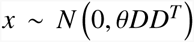 is a vector of tissue-specific random effects describing methylation effects and 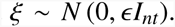. The matrices *J*, *A*, *B*, *C*, and *D* are design matrices with *B* being a function of genotype, *C* is a function of both genotype and methylation data and finally, *D* is a function of just the methylation data. *J* is *nt* × *t* dimensional matrix denoting the design matrix for the tissue-specific intercepts. *A* is *nt* × *nt* design matrix for the subject-specific intercepts. *B* is a *nt* × *t* design matrix of stacked genotypes. *C* is a *nt* × *t* design matrix of stacked (product of) tissue-specific methylation and genotype data. *D* is *nt* × *t* design matrices of stacked tissue-specific methylation data. The parameters of interest are *γ*, *δ*, *β* and *φ*; *α*, *λ*, *τ*, *θ* and ∈ are nuisance parameters. Alternatively, we can represent the distribution of *Y* conditional on methylation and genotype as –

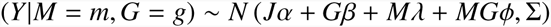

From our model, the log-likelihood function of the parameters conditional on the genotype and methylation data is given by–

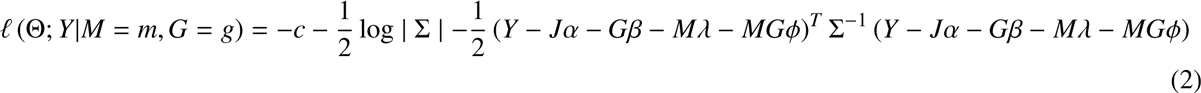

where Θ represents the vector of all the variance components involved in Σ and *c* is a constant. We test the null hypothesis that *H*_0_: *β* = *φ* = *γ* = *δ* = 0, i.e. the variant does not affect gene expression across any of the tissues. To do so, we compute the efficient scores for *γ*, *δ*, *β*, and *φ* by projecting off components correlated with the nuisance parameters. The reduced model under the null is –

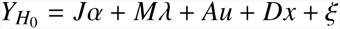

The efficient scores evaluated under the null are given by –

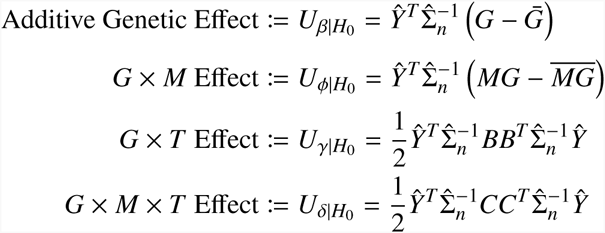

where 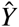 are the residuals from the model, 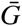 is an *nt*-dimensional vector of mean-centered genotypes,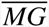 is an *nt*-dimensional vector of mean-centered product of genotypes and methylation, and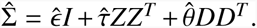. Our joint score test will test for the effect of genotype on 1) an overall shift in the gene expression, 2) tissue-specific interaction (*G* × *T*), 3) overall methylation (*G* × *M*), and 4) tissue-specific methylation (*G* × *M* × *T*). More on the individual components of our score test can be found in the supplementary section.

We propose a weighted sum of the above components (under the null) to arrive at our joint score test statistic, *U_ζ_*. Since *U_β_* and *U_φ_* are linear in *Y* while *U_γ_* and *U_δ_* are quadratic, we propose the following rule to combine them –

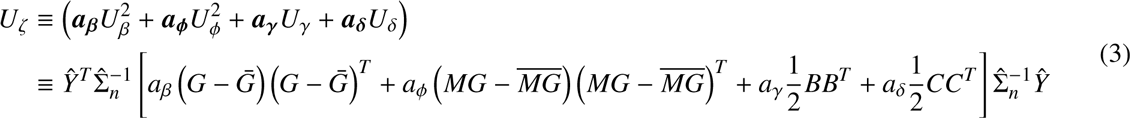

where *a_β_*, *a_φ_*, *a_γδ_* and *a_δ_* are scalar constants chosen to minimize the variance of *U_ζ_*. Under the null, *U_ζ_* is distributed as a mixture of chi-square random variables. We use Satterthwaite method [20] to approximate the *p* values from a scaled *χ*^2^ distribution by matching the first two moments as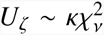 where 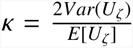 and 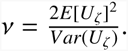

### Simulations

For a positive integer *t* that represents number of tissues, if **1** denotes a column vector of *t* ones and I denotes the corresponding *t* × *t* diagonal matrix, following the *t*-variate normal law denoted by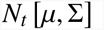 with mean *μ* ∈ R^*t*^ and variance Σ ∈ R^*t*^ ^×^ *^t^*, expression levels of a target gene *j* at a single locus by using the following vectorized form of the linear mixed model –

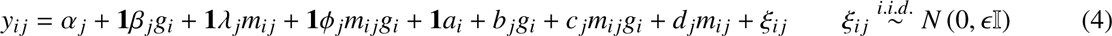

where *y_ij_* is a *t* × 1 vector of gene expression data, *α_t_* is the tissue-specific intercept (*α_t_* ∈ R^*t*^), φ *j* describes the main additive genotypic effect (φ _*j*_ ∈ R^1^), λ _j_ describes the overall effect due to methylation (*λ* _j_ ∈ R^1^), *φ* describes the interaction effect between the overall methylation and genotype (*ϕ* _*j*_ ∈ R^1^), *g_i_* is the value of a bi-allelic genotype such that *g* ∈ (0, 1, 2) represents the number of copies of the minor allele. The random effect *b_j_* ∈ R^*t*^ represents tissue-specific effect of the genotype, *c_j_* ∈ R^*t*^ represents tissue-specific interaction effect between methylation and genotype, *d_j_* ∈ R^*t*^ represents tissue-specific methylation effect, and a_i_ ∈ R^1^ is a subject-specific random intercept. We assume that all the random effects are independent and that *a_i_* ~ *N*_1_ (0, *τ*), *b_j_* ~ *N_t_* (0, *γ*I), *c_j_* ~ N_t_ (0, *δ*I) and *d_j_* ~ N_t_ (0, *θ*I). Methylation data for 5 tissues was generated independently from a multivariate normal distribution with mean zero and positive definite variance-covariance matrix.

We use 1,000 data replicates to evaluate the type I error and for power calculations. Simulations were performed by varying the following parameters-*β* (additive genetic effect), *φ* (*G* × *M* effect), the proportion of variation explained by the *G* × *T* effect 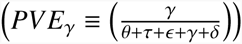 and the proportion of variation explained by the *G* × *M* × *T* effect 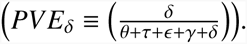 A linear mixed effects model was fit using the package *lme4*[21, 22] in the statistical environment R (R Core Team). The significance of an association between a mRNA - SNP pair in a tissue-by-tissue (TBT-eQTL) analysis is assessed by the *p* value obtained using *lm* function in R by fitting the following linear regression model.

For each mRNA - *cis*SNP pair, TBT-eQTL analysis was performed using the following linear regression model

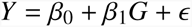

where *Y* is either gene expression data and *G* represents genotypes encoded as the number of copies of minor allele. The test statistic is the minimum *p* value over the total number of tissues from linear regressions performed separately in each tissue for each mRNA - SNP pair. Statistical significance was determined at a nominal *p* value of 0.05 for all power simulations (in case of TBT-eQTL analysis, it is 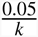 where *k* is the number of tissues).

## Preprocessing Gibbs et al datasets

### Data description

Fresh frozen tissue samples of the cerebellum (CRBLM), frontal cortex (FCTX), caudal pons (PONS) and temporal cortex (TCTX) were obtained from 150 neuropathologically normal samples [4]. Genotyping was performed using Infinium HumanHap550 beadchips (Illumina) to assay genotypes for 561,466 SNPs, from the cerebellum tissue samples. CpG methylation status was determined using HumanMethylation27 BeadChips (Illumina), which measure methylation at 27,578 CpG dinucleotides at 14,495 genes. Profiling of 22,184 mRNA transcripts was performed using HumanRef-8 Expression BeadChips (Illumina) The datasets are publicly available (GEO Accession Number: **GSE15745**; dbGAP Study Accession: **phs000249.v1.p1**).

### Gene expression data

Gene expression on four brain regions are publicly available as rank-invariant [23] normalized gene expression data (^“^series matrix file^”^). All the negative values in the gene expression dataset are changed to a 1 and the entire dataset was then log2 transformed. Before generating the PCA plots, samples with African and Asian ancestry were removed from the analysis. All the gene expression probes on sex chromosomes X and Y were removed from the analysis. In order to identify outliers in the PCA analysis, a simple yet standard approach or rule has been adopted. All the samples that did not follow the inter-quartile range (IQR) rule (*median* + 1.5 ∗ *IQR*) were excluded from further analysis (one CRBLM, one FCTX and two PONS). These samples were also eliminated by Gibbs *et al* in their original analysis.

Each gene expression probe was then adjusted for the biological and methodological covariates such as tissue bank, gender, hybridization batch and numeric covariates such as post-mortem interval (PMI) and age in order to remove any associated confounding effects using the following linear model –

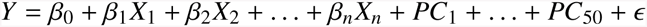

where Y is the gene expression data, *X*_1_…*X*_n_ represent the aforementioned biological covariates and systematic hybridization batch effects while *PC*_1_…*PC*_50_ are the top 50 principal components obtained from the original gene expression data. In order to target the difference in the genetic variation of expression among tissues, global variation in expression among tissues was removed by using the residual expression for each probe in each tissue after removing 50 PCs for further downstream analyses. It was shown in the past that the number of *cis-* eQTL detected significantly improved when 50 PCs were removed from the expression data [24].

### Methylation data

Methylation data, obtained as a ^“^series matrix file^”^ consisted of Beta-values, which represent the ratio of methylated probe intensity and the overall intensity (sum of methylated and unmethylated probe intensities) [25]. The methylation data was later adjusted for the biological and methodological covariates such as tissue bank, gender, hybridization batch and numeric covariates such as post-mortem interval (PMI) and age in order to remove any associated confounding effects using the following linear model –

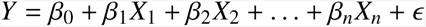

where Y is the methylation expression data and *X*_1_…*X*_n_ represent the aforementioned biological covariates and hybridization batch effects. The residual methylation expression was later used in the subsequent downstream analyses.

### Genotype data

The genotype data was obtained from dbGAP database (**phs000249.v1.p1**) following requisite author permissions. The genotype data was recoded into a SNP matrix of values 0, 1 and 2 representing minor allele counts. Samples with African and Asian ancestry were removed from the analysis. These SNPs were filtered on the missing-ness of the individual data and the SNP data (excluded SNPs with missing values), followed by MAF (included SNPs with MAF ≥ 0.05)and Hardy-Weinberg equilibrium (HWE; p-values ≤ 0.001) in the same order using PLINK [26] software. The resulting dataset has 400,097 SNPs after preprocessing.

## Results and Discussion

### Evaluating our new score test using Monte Carlo simulations

We evaluate our approach through extensive simulation studies. Briefly, each Monte Carlo simulated dataset was comprised of data from a single locus and a single gene, whose expression is measured across 5 tissues in 100 observations. For a given mRNA - SNP pair, the genotypes at each SNP in all the individuals were simulated as Binomial(2,0.3), i.e. a minor allele frequency 0.30 and assuming Hardy-Weinberg equilibrium. Methylation data for 5 tissues was generated independently from a multivariate normal distribution with a positive definite variance-covariance matrix. Since all the tissue-specific effects are modeled as random effects, a test of whether there are any tissue-specific effects is equivalent to testing whether the variances of the random effects (*γ* and *δ*) are zero. Thus, our model involves testing four scalar parameters (*β*, *φ*, *γ* and *δ*). Simulations under the null hypothesis confirm that our method has the correct type 1 error (see supplementary material). Since we model the effects of both epigenetic and genetic variation, we evaluated any power loss in identifying mRNA - SNP associations in the absence of any epigenetic effect. This was accomplished by comparing our method’s performance with TBT-eQTL approach by keeping all the parameters associated with methylation in equation 1 at zero (i.e. *λ* = *φ* = *δ* = *θ* = 0). We also compared our method with a previously proposed multi-tissue eQTL method, implemented in our software JAGUAR [**?**], which is made available at Comprehensive R Archive Network (CRAN) repository. Briefly, JAGUAR implements an approach that jointly models the overall shift in the gene expression due to genotype together with tissue-specific interaction with genotype in order to efficiently identify multi-tissue eQTL. From Figure 2a, we see that JAGUAR outperforms both TBT-eQTL and our new joint score test. This loss of power, though not substantial, may be attributed to testing for an inexistent methylation effect. However, in the presence of a methylation effect our method outperforms both TBT-eQTL and JAGUAR as evidenced by Figure 2b.

**Figure 2.**
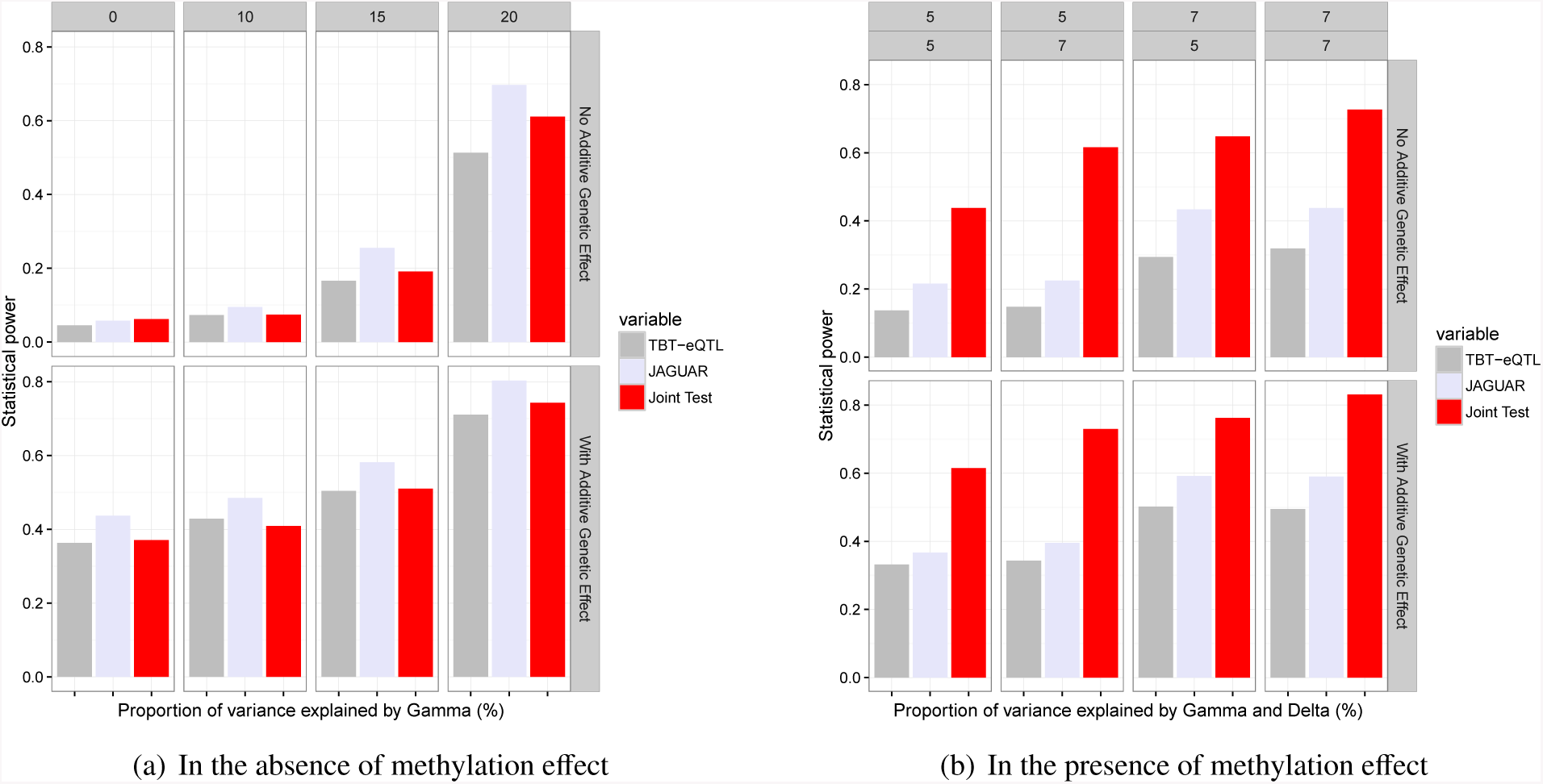
Mapping eQTLs in multiple tissues using 1BT-eQTL, JAGUAR and Joint Score Test methods. Left panel illustrates eQTL mapping in the absence of higher order methylation effects. Right panel illustrates QTL mapping in the presence of higher order methylation effects and the numbers in the top two rows indicate the proportions of variance explained by both *γ* and *δ*, respectively.

We also compared our joint score test to a TBT-eQTL approach that included methylation as a baseline covariate [16], henceforth referred to as TBTm-eQTL analysis, using the following linear regression model –

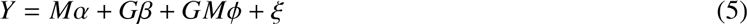

where *Y* is a *nt*-dimensional matrix of expression levels in *t* tissues and *n* individuals, α is a fixed effect representing the tissue-specific intercepts, *G* is a *nt*-dimensional matrix of genotypes, *β* is a fixed effect of genotype across all tissues, *M* is a *nt*-dimensional matrix of methylation information and *φ* is genotype × methylation interaction effect (fixed effect). Minimum *p* value from the TBTm-eQTL analysis across all the tissues is computed for power calculations. Table 1 shows that out method significantly outperforms 1BTm-eQTL approach owing a clear statistical advantage in using our joint score test over the TBTm-eQTL approach.

**Table 1:**
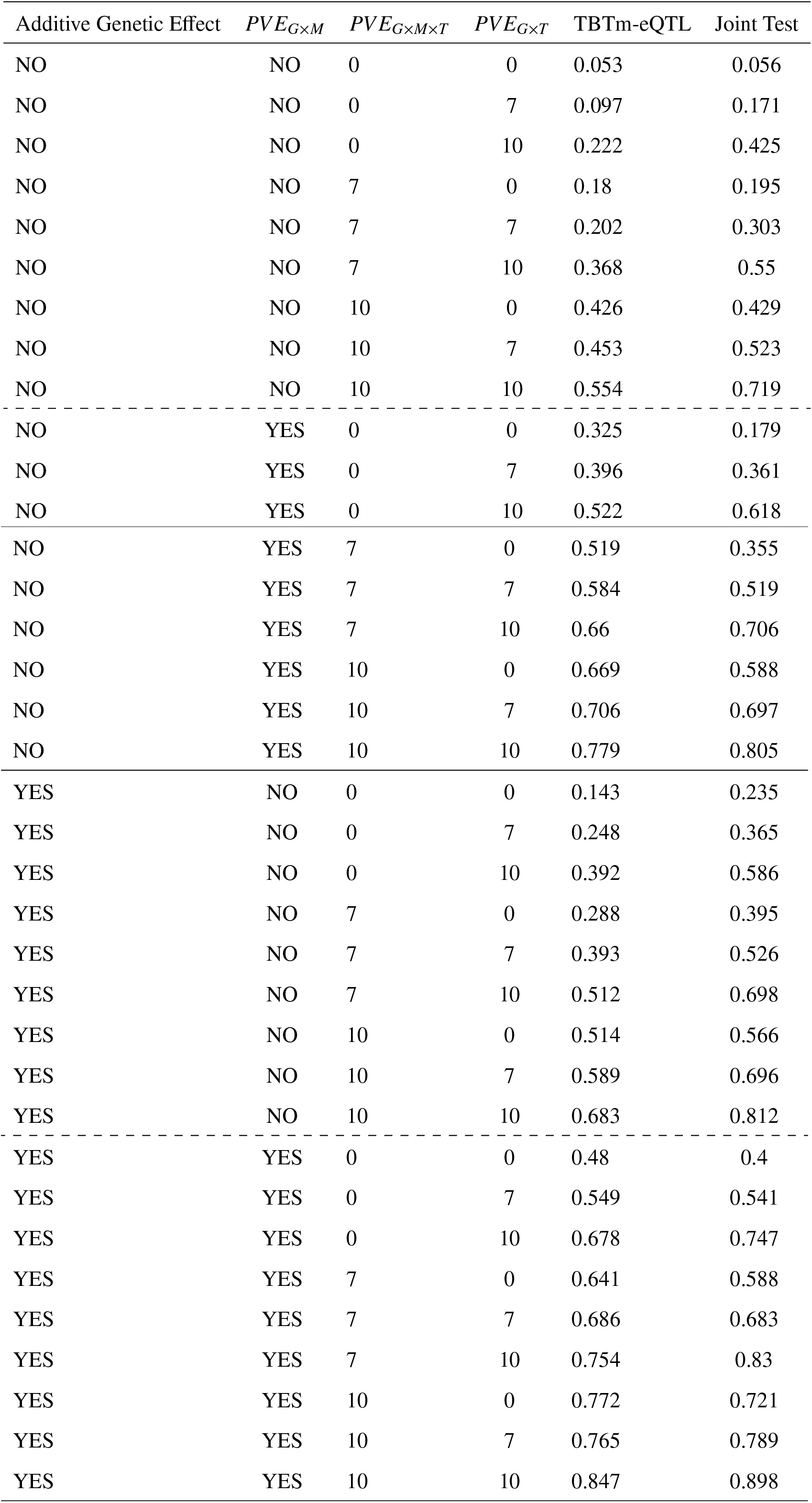
Table comparing the statistical power of our method and TBTm-eQTL approach. This data were generated from 1,000 simulations run on 100 individuals and five tissues with genotypes generated at a common variant allele frequency (MAF = 0.3).

### Region-specific DNA methylation impacts eQTL mapping in adult human brains

In order to demonstrate the effectiveness of our method, we applied it to Gibbs *et al* [4] dataset comprising of 150 individual data obtained from four regions of human brain. We performed data analyses that focused on only *cis* candidate regions. The proximity of an eQTL to the transcription start site of a gene did not exceed 100 kilobase up-and down-stream of the transcription start site of a gene/mRNA transcript (*cis*-SNP). CpG islands that were less than 1.5 kilobase up- and down-stream of the transcription start site of the same mRNA transcript were paired with the mRNA transcripts. Figure 3 illustrates the analysis design in one tissue. Each mRNA transcript was tested for an association with every *cis*-SNP in the presence of a (methylated or unmethylated) CpG site located in the promoter region.

**Figure 3.**
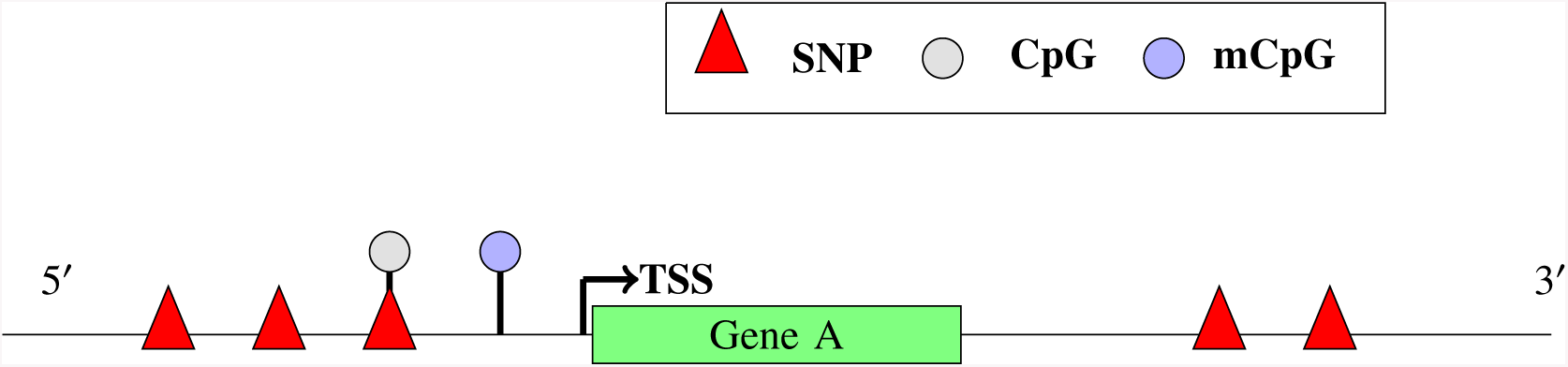
Here is an illustration of the analysis design using a hypothetical Gene A. The red triangles indicate SNPs and the circles, CpG sites (gray = unmethylated; blue = methylated). CpG sites that are at least 1.5 Kb from the transcription start site (TSS) of a gene were picked for the analysis. All the SNPs that were picked did not exceed 100 kilobase up- and down-stream of the transcription start site of a gene (*cis*-SNPs).

Our joint score test method performed a total of 491,496 tests (totaling 11,095 mRNA transcripts, 14,663 CpG sites and 144,571 *cis*SNPs). Each mRNA transcript may have multiple CpG sites in its promoter region. Thus, each such mRNA - CpG pair is tested for an association with a *cis*SNP. It is important to note that our method does not test any direct association between an mRNA transcript and its corresponding CpG site. Any resulting combinations of mRNA transcript, CpG site and a SNP would describe the relationship between the mRNA and SNP in the presence of the corresponding promoter CpG site. Our method identified a total of 13,212 such combinations corresponding to 9,065 eQTLs that are statistically significant at 5% false discovery rate (FDR). In order to account for the number of traits being tested, the *p* values obtained from applying our joint score test were adjusted for multiple testing using an optimized FDR approach to obtain *q* values (FDR adjusted *p* values) [27]. We observed that majority of these significant results are driven by a combination of additive genetic effect (86%) and *G* × *T*effect (43%) while the *G* × *M* and *G* × *M* × *T* effects were barely observed. This may be due to a lack of any distinct tissue-specificity in the methylation data, which we observed while preprocessing Gibbs *et al* data (see Methods section). However, we expect that the aforementioned effects may be well pronounced across diverse cell-types such as the ones made available by the Genotype-Tissue Expression Project (GTEx) [28].

We performed two region-by-region or TBT approaches on the same set of mRNA transcripts, CpG sites and SNPs as above, one with DNA methylation as a covariate (TBTm-eQTL) and the other with no methylation (TBT-eQTL) and compared the results with our approach. We estimated *q* values from each set of *p* values (originated from each region-by-region analysis) and minimum *q* value for a given mRNA - SNP pair across all the brain regions was computed, which indicates the presence of a statistically significant pair in at least one brain region. The number of significant associations in at least one brain region were then assessed at 5% FDR (*q* value 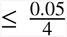 where 4 is the number of brain regions). TBT-eQTL approach identified a total of 11,014 mRNA-*cis*SNP pairs significant in at least one region of the brain at 5% FDR. Roughly 79% of these TBT-eQTLs overlap with eQTLs identified using our method. On the other hand, TBTm-eQTL approach identified 10,926 combinations of RNA transcripts, CpG sites and SNPs corresponding to 7,496 eQTLs with a 73% overlap with eQTLs identified using our method.

In order to assess the role of brain region-specificity on gene expression and the advantages in jointly modeling all the brain regions on mapping eQTLs, we compared our joint score test approach with a previously proposed multi-tissue eQTL mapping method [19] implemented by our software JAGUAR. JAGUAR identifies 16,919 eQTLs (96% of them overlap with the TBT-eQTLs, 80% of them overlap with TBTm-eQTLs, and 92% of them overlap with the joint tests’s eQTLs) at 5% FDR. All the eQTLs that overlap between JAGUAR and our new joint score test are mostly driven by the additive genetic effect and *G* × *T* effect and not higher order methylation interaction effects such as *G* × *M* and *G* × *M* × *T*. This absence of any pronounced region-specific DNA methylation effect explains the lower number of eQTLs identified by our method. However, as we have shown using simulation data, in the presence of any region-specific interaction effects involving methylation, our joint score test is far more informative than the results from JAGUAR. There are a total of 744 eQTLs identified by our method that weren’t found to be statistically significant by JAGUAR. This is because 90% of these eQTLs were driven by *G* × *M* × *T* interaction effect, which is not tested by JAGUAR. For example, let us consider a protein coding gene Transmembrane BAX inhibitor (TMBIM1; Ensemble ID - ENSG00000135926), located on chromosome 2, which has 18 annotated SNPs (possibly in LD with each other) and two promoter CpG sites in our preprocessed datasets. Out of these 36 (number of mRNA - CpG pairs × the number of SNPs) combinations of mRNA transcript, CpG sites and SNPs and a possible 18 eQTLs, our method identified 13 to be statistically significant. None of the 36 eQTLs were found to be statistically significant by any TBT-based or the multi-tissue eQTL approaches (Figure 4a). Upon close inspection, we observed that there is an absence of any additive genetic effect or a *G* × *T* interaction effect however, the presence of *G* × *M* × *T* interaction effect is driving the statistical significance. This is a good example of mapping eQTLs by leveraging effects due to DNA methylation. Of note, TMBIM1 is ubiquitously expressed in brain [29] and BAX-inhibiting peptides have been known to prevent neuronal cell-death induced by oligomeric β-amyloid, which plays an important role in the pathogenesis of Alzheimer disease [30]. Out of the 13 significant eQTLs for TMBIM1 that our method identified, Figure 4b illustrates the association between the gene, top CpG site (Methylation Probe ID:cg14849559) and SNP (SNP ID: rs921970) with the lowest *q* value (*q* val = 0.00386).

**Figure 4:**
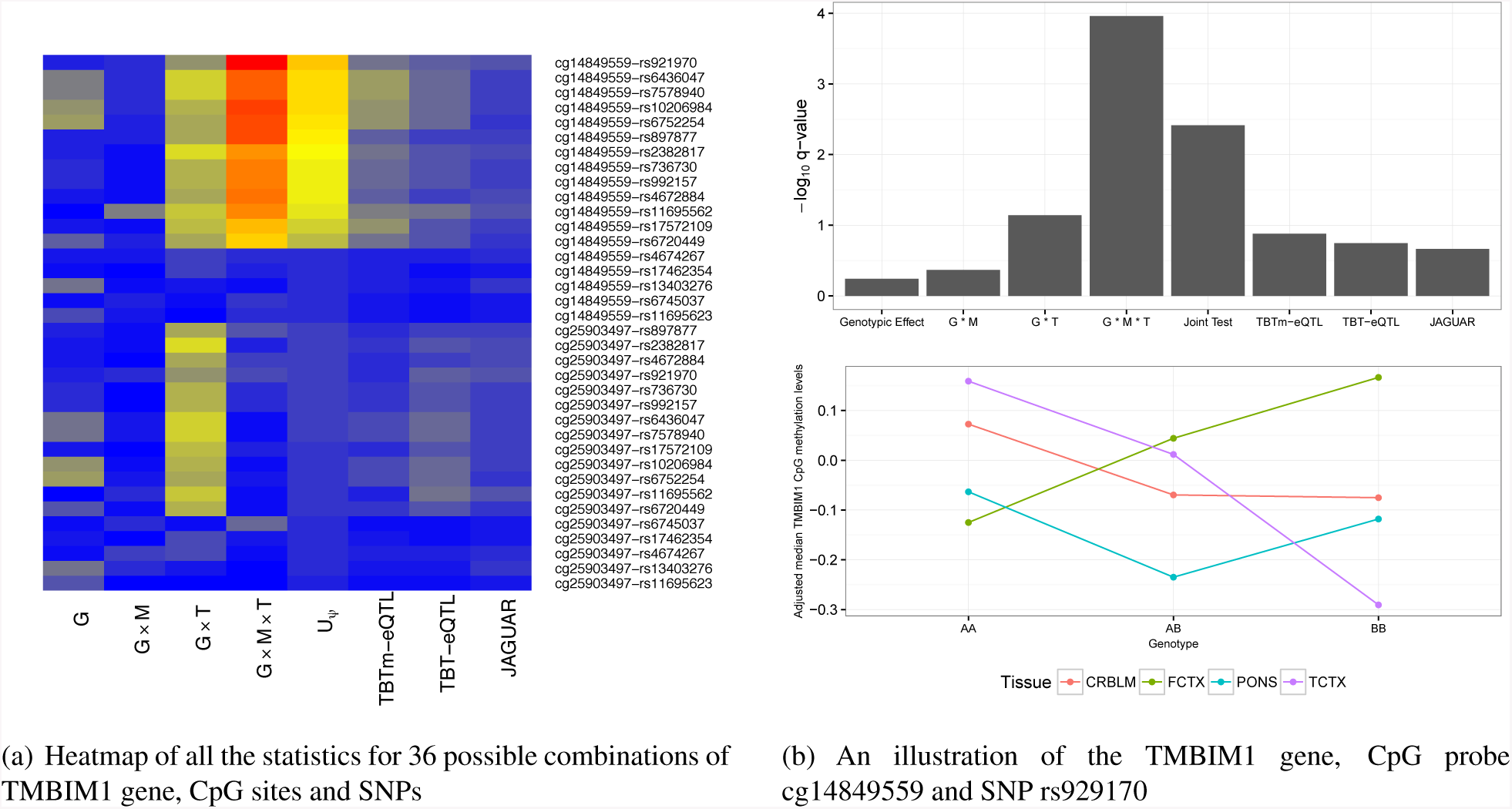
Figure 4a indicates the heatmap of –*log*_10_ *q* values of all 36 possible combinations of TMBIM1 gene,tawo CpG sites and 18 SNPs. Red in the heatmap indicates a lower *q* value and thus higher statistical significance where as blue indicates otherwise. Figure 4b has two panels. Top panel displays all the statistics computed for TMBIM1 gene, CpG probe cg14849559 and SNP rs929170. The first four bars indicate the four different effects tested by our joint score test and 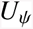 represents the omnibus *q*value of our joint test. Bottom panel illustrate the *G* × *M* × *T* interaction plot.

On the other hand, we also see many instances of eQTLs that were observed to be statistically significant using JAGUAR but not our joint score test method due to the lack of any DNA methylation effects. For example, JAGUAR identified gene B-Cell CLL/Lymphoma 2 (BCL2; Ensembl ID - ENSG00000171791), a gene that promotes neuronal cell death or apoptosis [31], to have a statistically significant association with a promoter eQTL (SNP ID: rs17676919), as illustrated by Figure 5. As seen in this figure (from –*log*_10_ *q* values), the lack of any higher order methylation effects may have resulted in not being identified as a potential eQTL by our joint score test method.

**Figure 5:**
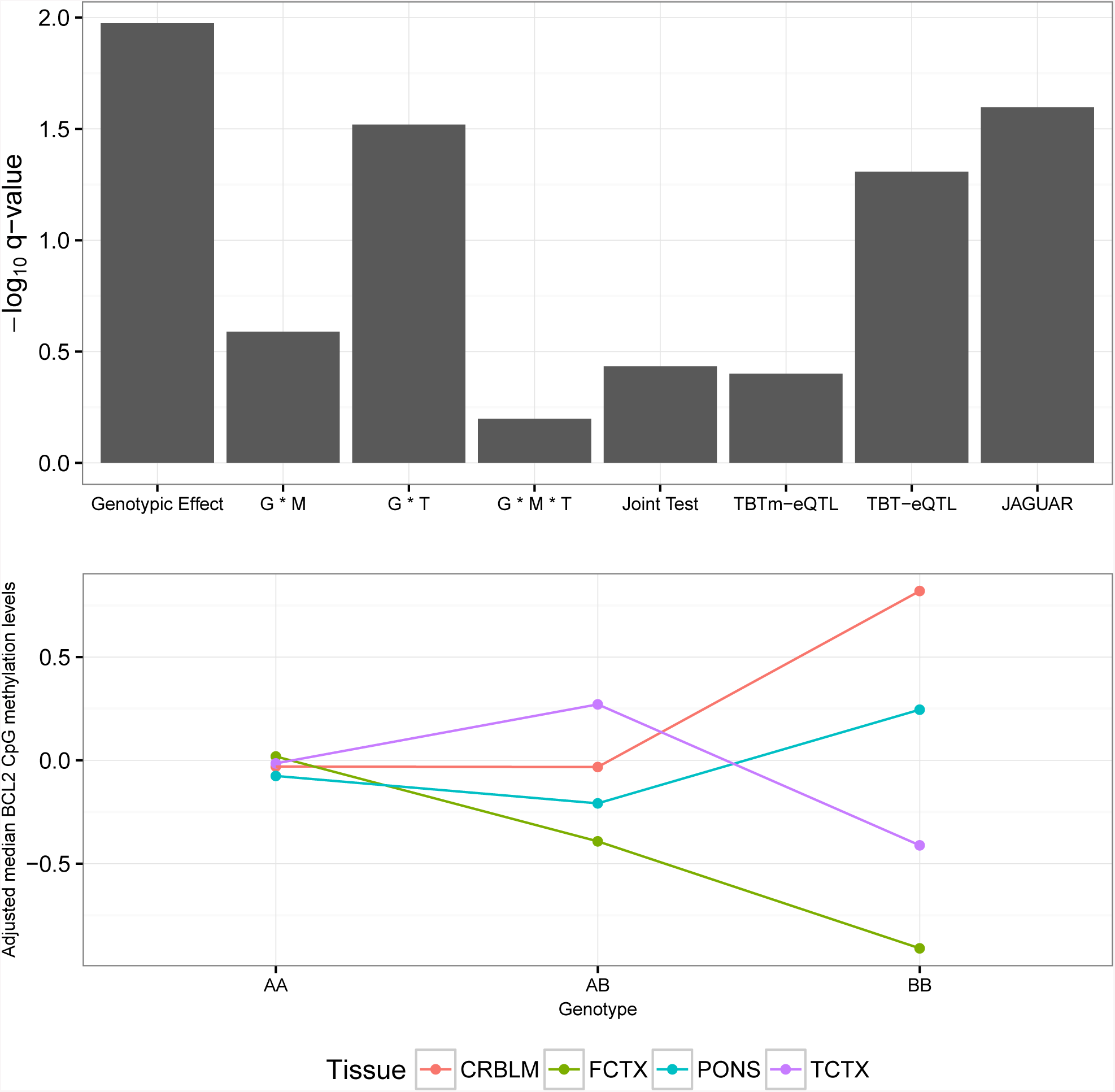
An example of an eQTL for gene BCL2 not identified as statistically significant by our joint score test method. Top panel displays all the statistics computed for BCL2 gene, CpG probe cg14455307 and SNP rs17676919. The first four bars indicate the four different effects tested by our joint score test and 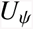 represents the omnibus *q* value of our joint score test. Bottom panel illustrate the G × *M* × *T* interaction plot.

To assess the biological relevance of the genes with eQTLs identified by TBT or multi-tissue methods including our new joint score test, we performed a KEGG pathway term enrichment analysis [32] for each set of results separately (see Supplementary materials). KEGG pathways were considered overrepresented if a set of at least three genes from different linked regions is observed to be overrepresented with an adjusted significance level of q value < 0.05, calculated from a hypergeometric test. Our method identified 5 overrepresented pathways (Metabolic pathways, Ribosome, Fatty acid degradation, Purine and Pyramidine metabolism), JAGUAR identified 2 pathways while TBT-eQTL identified 1 overrepresented pathway. The overrepresented pathway, ^“^Metabolic Pathways^”^ (KEGGID: hsa01100) is the only common pathway between TBT-eQTL, JAGUAR and our method methods. On the basis of prior knowledge of function, the overrepresented pathways ^“^Purine metabolism^”^ (KEGGID: hsa00230) and “Pyramidine metabolism” (KEGGID: hsa00240) are plausible functional candidate pathways for schizophrenia [33]. These information can be used to guide genetic analyses by selecting these relevant pathways and genes associated with the pathways for schizophrenia.

## Conclusion

Overall, our efforts are primarily directed to understanding two very specific aspects – 1) the overall effect of a genetic variant on gene expression regulation by accounting for any changes in tissue DNA methylation levels, and 2) map eQTLs by leveraging tissue-specific methylation effects. Currently, there are no methods that jointly model the epigenetic and genetic control of tissue-specific gene expression. Many eQTL studies fail to account for the masking effect on a genetic variant due to DNA methylation, which may regulate gene expression across multiple tissues. Our method provides an efficient framework to integrate SNPs, DNA methylation and gene expression, and investigate how the different forms of variation inter-relate.

The dataset examined here used genome-wide association (GWA) study SNP array platform to interrogate germline variation that includes an overwhelming number of common variants. Although GWA studies have been able to explain a small fraction of the genetic components of common human diseases, it is hypothesized that some of the missing heritability may be due to rare variation. Since standard common disease common variant approaches are severely underpowered to tease out any underlying variants that are moderate to extremely rare, there is an emphasis on large sample sizes and gene-based association tests in order to securely identify genetic risk factors that may otherwise be outside the range detectable by GWA studies [34]. One solution to the aforementioned issue would be to prioritize genetic variants in a non *ad-hoc* framework that preferentially weights genetic variants. Our method can provide a statistically disciplined weighting framework within which genetic variants can be either up- or down-weighted for any subsequent downstream analyses. Our method may also be useful in generating weights to any methods that use a reference data set in which both genome variation and gene expression levels have been measured to develop prediction models for gene expression [35].

Since we are modeling the effects of non-coding regions (via CpG sites) on gene expression using our model, we can easily use micro-RNA (miRNA) data instead of CpG site methylation data and model post-transcriptional regulation of tissue-specific gene expression. miRNA expression, also a tissue-specific phenomenon, have been known to post-transcriptionally silence expression of mRNA transcripts. The presence of genetic variants such as SNPs may have an effect on the biogenesis and function of miRNA molecules leading to a downstream effect on gene expression [36]. This tissue-specific interaction between miRNA and SNP can be modeled in a similar fashion, analogous to modeling the interaction effects of tissue-specific DNA methylation and SNPs. The flexibility of our model also enables us to incorporate new information such as gene isoform data and accommodate the analysis of next-generation sequencing data (such as RNA-seq) by modeling gene transcripts in an analogous fashion to tissues in our current model formulation. This type of analysis would aggregate expression over all the splice variants of a gene across multiple tissues and inform us of tissue-specific alternative splice variant of a gene. These results become relevant to studying genetic effects on alternative splicing and its key role in important cellular networks.

